# Network and structural analysis of quail mucins with expression pattern of MUC1 and MUC4 in the intestines of the Iraqi Common Quail (Coturnix Coturnix)

**DOI:** 10.1101/2023.07.18.549497

**Authors:** Hazem Almhanna, Aqeel Mohsin Mahdi AL-Mahmodi, Abdulrazzaq B Kadhim, Arun HS Kumar

## Abstract

**Background:** Mucins have vital pathophysiological role in gastrointestinal tract (GIT) of avian and other species. However, despite this very little is known about the types of mucins expressed in quail GIT. Hence in this study we examined the expression pattern of mucins (MUC1, and MUC4) in the GIT of the Iraqi Common Quail (Coturnix Coturnix) and performed the network and structural analysis of all reported types of mucins in various breeds of quails.

**Materials and methods:** This study protocol was approved by the animal ethics research committee of the College of Veterinary Medicine, University of Al-Qadisiyah, Iraq. Fresh samples of small and large intestines were used for histological and gene expression analysis of MUC1, and MUC4. Network and structural analysis of all reported types of mucins in quails was performed using the STRING Database, Chimera software and PrankWeb-Ligand Binding Site Prediction tool.

**Results:** The histological analysis using Alcian blue and PAS stains indicated that most mucins in the intestines of quails were of the acidic mucin type, with minimal prevalence of neutral mucins. The expression of acidic mucins was relatively higher in the duodenum, ileum, caecum, and colon, while the jejunum showed a relatively higher expression of neutral mucins. Gene expression analysis revealed higher expression levels of MUC1 and MUC4 mRNA in the jejunum and colon, with its least expression in the duodenum and ilium. Network analysis indicated predominantly mucin-mucin interactions, with MUC 1, 15, 16 and 24 showing preferential homologous networks while the MUC 2, 4, 5 and 6 showed heterologous networks. Detailed evaluation of intermolecular hydrogen bond formation highlighted the interactions between specific mucin combinations, with certain combinations showing higher affinity, such as MUC5A-MUC6, MUC5A-MUC5B, and MUC5B-MUC6. In contrast, MUC15, MUC16, and MUC24 exhibited limited interactions with other mucin types. Binding site analysis indicated that MUC5B and MUC6 had the most number of binding sites with high probability scores, while MUC2, MUC4, and MUC5A showed lower probability scores despite having more binding sites. In contrast MUC 1, 15, and 16 had very few binding sites (<3 binding sites) all with very low probability scores.

**Conclusion:** The findings of this study provide valuable insights into the composition, expression, network interactions, and binding sites of mucins in the quails, contributing to the understanding of mucin-related processes in gastrointestinal physiology and potential implications for gastrointestinal diseases.

## Introduction

Mucins are glycoconjugates proteins which include many members, and have highly O-glycosylated glycoproteins[1]. Mucins can be found in two forms i.e., 1) as membrane associated proteins which project from cell surface against extracellular agents and 2) as soluble form proteins which can be detached and form gel of the mucus[2]. Mucins have large molecular weight, and consist of tandem repeat structure which are mostly rich with serine and threonine backbone varying in length depending on the type of mucins[3]. Different types of mucins interact with various biomolecules including monosaccharides, polysaccharides, and proteoglycans and regulate many functions [4, 5]. Also, mucins constitute the main component of the mucus and together with soluble mucin[6, 7] play a vital role in the defence mechanism of the intestine by forming a mucosal barrier covering the intestinal tract and prevent adherence of infectious agents to the intestinal epithelium[8-10].

There are many pathogens that affect the digestive system of the birds including parasites and microbial infections which cause pathological changes to the different parts of gastrointestinal tract. These diseases might be caused by bacterial infections for example salmonella species, and E. coli [11, 12] or protozoan infestations such as giardia, and trichomonads, and coccidia[13, 14], or parasites such as tapeworms, and roundworms[15, 16]. Mucins in gastrointestinal tract are essential for the defence against these pathogens through both humoral immunity and non-specific defence mechanisms[17]. Specifically MUC1 and MUC4 which are highly O-glycosylated glycoproteins, have large molecular weight, are membrane bound [18, 19] are reported to be highly expressed in the colon of the human [20] and gastrointestinal tract of pig [21]. The differential expression and molecular changes of MUC1 and MUC4 are reported to be useful prognostic biomarker in many cancers including prostate cancer, mucoepidermoid carcinomas of salivary glands, pancreatic cancer, hyperplastic polyps, serrated adenomas, breast cancer, gastric carcinoma and traditional adenomas of the colorectal cancer [20, 22-25]. Additionally the expression and signalling pathways of MUC1 and MUC4 have been associated with oncogenic role and responsible for initiation and progression of the tumours in different locations of the body[26, 27]. Despite the well-recognised role for mucins in the quail GIT, data on detailed expression of different mucins in quail gastrointestinal tract are lacking. Hence in this study expression of MUC1 and MUC4 in different parts of small and large intestine of the Iraqi common quail (Coturnix Coturnix) was examined and a detailed network and structural analysis of all reported types of mucins in various breeds of quails were examined to assess their biological significance.

### Material and Methods Samples and study design

Ten Iraqi common quail birds were used for this study, birds were healthy and killed following animal ethics rules of College of the Veterinary Medicine-University of Al-Qadisiyah, Iraq (Approval P.G. No. 1890). Quails were euthanised by cervical dislocation and the abdomen was dissected to collect specimens of the duodenum, jejunum, ilium, caecum, and colon for histology and gene expression analysis. Specimens were separated into two groups, specimens saved in TRIZOLe (SRCr Green-Zol reagent. Iraq) for Quantitative Real-Time PCR (qPCR) study and other specimens saved in 10% neutral buffered formalin (100ml formalin (37-40% stock solution), 900ml distilled water, 9g of NaCl, and 12g of Na2HPO4 (dibasic/anhydrous) for routine and special stains study.

### Histological analysis

The specimens of quails collected for histology analysis were fixed in 10% neutral buffered formalin and left for 48 hours before histological processing. The tissue sections were stained by Haematoxylin and Eosin (H&E) and Periodic acid–Schiff (PAS) stain combination with alcian blue stain as previously described [28, 29]. All slides were examined using a light microscope (Olympus, Japan), and representative images from different regions of the tissue were captured at following magnification (4x,10x, 20x, and 40x).

### RNA Extraction and cDNA synthesis

Total mRNA was extracted from small and large intestine of quails using the Accuzol® reagent kit (Bioneer. Korea) as per the manufacturers instructions. Briefly, 200 mg of tissue was weighed for each part of intestine in a 1.5ml Eppendorf tube, then 200ul of chloroform was added and mixed, and incubated on ice for 5 minutes. Next, tissues were centrifuged at 12000 rpm, 4C°, for 15 minutes, and the supernatant was collected. After that, 500ul of isopropanol was added and mixed, and incubated again in cooled conditions (4°C) for 10 minutes. Following this the samples were centrifuged at 12000 rpm, 4C° for 10 minutes. The supernatant was neglected, and 1ml 80% ethanol was added, mixed by vortex, and spun at 12000 rpm, 4C° for 10 minutes. The supernatant was discarded and pellet was stored in the Eppendorf tubes to air dry. To finish, 50ul of DEPC water was added to pellets of RNA to dissolve and stored at -20^0^C until analysis. Nanodrop spectrophotometer (THERMO. USA) was used to measure RNA concentrations for each samples. All samples were processed using the DNase I enzyme kit (Promega Company, USA) as per manufacturer’s instructions. Then, DiaStar™ OneStep RT-PCR Kit (China) was used to translate total RNA into cDNA following instructions and thermocycler conditions, after that, cDNA concentrations were normalised for all samples and saved at -20^0^C for next step.

### Quantitative Real-Time PCR (RT-qPCR)

RT-qPCR technique was applied for identifying the levels of MUC1, and MUC4 gene expression using the Real-Time PCR system (BioRad./ USA). Following primers were used in this study; Coturnix japonica glyceraldehyde-3-phosphate dehydrogenase (gene code: XM_015873412.2) (GAPDH housekeeping gene), forward primer: TGCTGGCATTGCACTGAATG, Reverse: CACGGTTGCTGTATCCAAACTC, and Coturnix japonica MUC1-like (LOC107317569) (MUC1), mRNA (gene code: XM_032448780.1) forward primer: TAATGCTGCCCCAATTGCTG, reverse primer: TGAGGTTGTATCCCAGTGCAG.

Coturnix japonica mucin 4, cell surface associated (MUC4), mRNA, code: XM_032446547.1) forward primer: AATGCAAAGTGCCACAGCTG, reverse primer: TTGGTGTTCCTCCAAAACGC. The amplification and normalization of GAPDH housekeeping gene, MUC1, and MUC4 genes expression used the SYBER Green dye qPCR master mix to detect the level of genes expression as per the kit instructions (AccuPowerTM 2XGreen Star qPCR master mix kit. Bioneer. Korea). The thermocycler conditions were set as the follows: initial denaturation was fixed at 50^0^C for hour, and repeated cycle one only, denaturation was at 95^0^C for 20 seconds, annealing\extension detection(scan) was at 60^0^C for 30 seconds and again repeated cycles at 45 seconds, and to end with melting temperature was at 60-95°C, for 0.5 seconds and repeated cycle one only.

### Network and binding site analysis of quail mucins

All the reported mucin sequences in quails were identified in the Uniprot database and their 3D structure were generated using the AlphaFold or Swiss homology modelling tools as reported previously. The network protein analysis of quail mucins was conducted as reported before using the STRING Database (https://string-db.org), to observe its functional protein-protein interactions. The 3D structure of the quail mucin’s in PDB format were imported onto the Chimera software and the number of hydrogen bonds (H-bond) formed between them at 10 Armstrong (10A) distance was evaluated. A heatmap of the number of H-bonds formed between different mucin combinations were generated to identify the high affinity interactions. The binding sites of each of the quail mucin’s were identified using the PrankWeb: Ligand Binding Site Prediction tool (https://prankweb.cz/).

### Statistical Analysis

The MUC1 and MUC4 gene expression data were calculated using RT-q PCR and the 2^ΔCT method [30, 31], and statistically, one-way ANOVA was applied for analysing RT-qPCR data using SPSS (IBM SPSS Statistics 23.0) and significance was considered at the P≤ 0.05.

## Results

### Histological assessment of quail intestines

H & E stains showed the general histological features of the duodenum, jejunum, ileum, caecum and colon of quail, and displayed four main layers involving mucosa, submucosa, muscularis, and serosa layers. The special patterns and structures of each part of the quail intestinal tract was consistent with avian literature. All the regions of quail intestinal tract showed three distinct layers of mucosa i.e., 1) simple columnar epithelium with goblet cells on basement membrane, 2) extended lamina propria with intestinal glands and 3) varying aggregations and centres of lymphocytes surrounded by loose connective tissue. The submucosa consisted of varying degree of intestinal glands in different regions of the intestine with fat globules, nerves, lymphatics and blood vessels. The muscularis region showed two layers, i.e., outer longitudinal and inner circular layer, which were externally covered by thin serosa layer (Figure 1).

**Figure 1:**
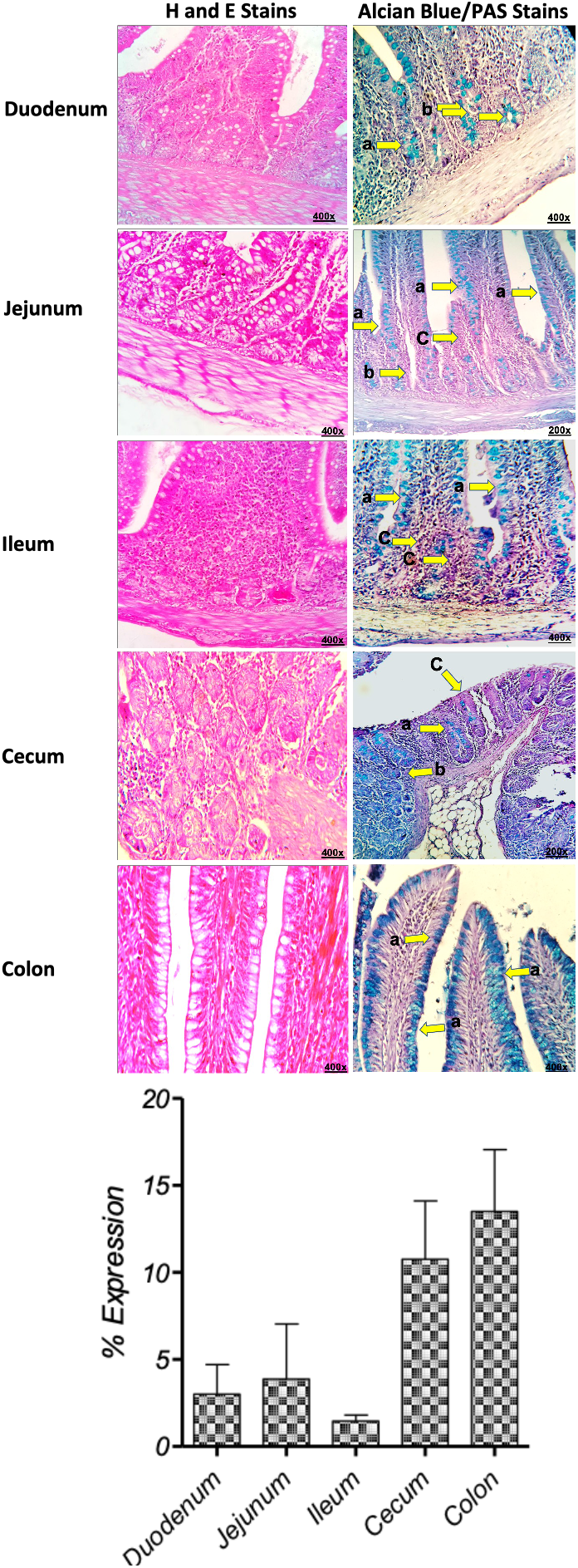
Histology of quail intestines. Representative histology sections of different parts of the small and large intestine of quail stained with H & E, and combination Alcian blue/PAS stains are shown. Photographs are labelled with a: epithelial tissue; and b: intestinal glands were positive stained with alcian blue, while c: epithelial tissue were stained with PAS stain (400x magnificence). The bar graphs show the semi-quantification of the alcian blue staining area. Data is shown as mean+/-SD of percent expression of alcian blue staining area in the histological section assessed in three independent histological section per tissue type.

The combination of Alcian blue and PAS stains allows the simultaneous visualization of both acidic and neutral mucins. Acidic mucins stain blue with Alcian blue, while neutral mucins and glycogen stain pink/magenta with the PAS reaction. This staining combination provides a comprehensive assessment of mucin composition within tissues. Most of the mucins of epithelium, goblet cells and cells of the intestinal glands of the duodenum, jejunum, ilium, caecum, and colon were stained blue colour (Alcian blue) with PAS stain being very sporadically visible. Hence the staining pattern observed in this study suggest that most mucins in the small and large intestine of quails are of acidic mucin type with very little prevalence of neutral mucins. Especially the expression of acidic mucins was relatively higher in duodenum, ilium, caecum and colon compared to that in jejunum wherein relatively higher expression of neutral mucins was observed (Figure 1). The semi-quantification of alcian blue staining suggested higher relative expression of acidic mucins in caecum, and colon compared to duodenum, jejunum and ilium (Figure 1). The expression of acidic mucins was observed to be least in the ilium. The PAS staining intensity was very light in the duodenum, and colon suggesting poor expression of neutral mucins in this part of quail intestine.

### Expression of selected mucin transcripts in quail intestines

In our study we specifically examined for the presence of MUC1 and 4 transcripts using RT-qPCR in various regions of the quail intestine. The amplifications and melting peaks showed consistent curve without any nonspecific product or amplification, with melting peak ranging from 80 to 88°C (figure 2) thus providing a validation for our method used. RT-qPCR amplification plots of MUC1 and 4 gene of the duodenum, jejunum, ileum, caecum, and colon were accurately detected and threshold cycles (Ct) numbers of expression of MUC1, and MUC4 were clearly noticed and ranged between CT 21.99 to CT 26.45 (figure 2, tables 1, and 2). The RT-qPCR amplifications of MUC1, and MUC4 mRNA was obviously determined in both small and large intestine (figures 2 and tables 1, and 2). The higher expression of MUC1 and MUC4 mRNA was observed in jejunum and colon while the expression of MUC1 mRNA was least in the duodenum and the expression of MUC4 mRNA was least in the duodenum and ilium (figure 2). Relatively higher expression of MUC1 was observed in jejunum and colon compared to MCU4 expression. While relatively higher expression of MUC4 was observed in duodenum and caecum compared to MUC1 expression.

**Table 1:** This table presented values of gene expression of housekeeping gene and MUC1 which were analysed using 2^ΔCT method.

| No. | Organs | CT (MUC1) | CT (GAPDH) | $\Delta CT$ | Fold change ($2^{\Delta\Delta CT}$) | Mean |
| --- | --- | --- | --- | --- | --- | --- |
| 1 | Caecum | 26.45 | 26.62 | 0.17 | 1.13 | 2.28 |
| 2 | Caecum | 24.72 | 26.61 | 1.89 | 3.72 |  |
| 3 | Caecum | 25.14 | 26.13 | 0.99 | 1.98 |  |
| 4 | Colon | 24.17 | 26.05 | 1.88 | 3.69 | 7.48 |
| 5 | Colon | 22.30 | 26.09 | 3.79 | 13.82 |  |
| 6 | Colon | 24.34 | 26.64 | 2.30 | 4.92 |  |
| 7 | Duodenum | 25.59 | 26.63 | 1.04 | 2.05 | 1.55 |
| 8 | Duodenum | 25.63 | 26.56 | 0.93 | 1.91 |  |
| 9 | Duodenum | 26.74 | 26.21 | -0.53 | 0.69 |  |
| 10 | Ileum | 21.99 | 24.63 | 2.64 | 6.25 | 3.80 |
| 11 | Ileum | 25.11 | 26.92 | 1.81 | 3.50 |  |
| 12 | Ileum | 24.83 | 25.56 | 0.73 | 1.65 |  |
| 13 | Jejunum | 23.54 | 27.11 | 3.57 | 11.87 | 7.31 |
| 14 | Jejunum | 23.00 | 24.45 | 1.45 | 2.74 |  |
| 15 | Jejunum | 22.96 | 25.83 | 2.87 | 7.32 |  |

**Table 2:** This table presented values of gene expression of housekeeping gene and MUC4 which were analysed using 2^ΔCT method.

| No. | Organs | CT (Mucin4) | CT (GAPDH) | $\Delta CT$ | Fold change ($2^{\Delta\Delta CT}$) | Mean |
| --- | --- | --- | --- | --- | --- | --- |
| 1 | Caecum | 26.86 | 26.62 | -0.24 | 0.84 | 2.59 |
| 2 | Caecum | 25.83 | 26.61 | 0.78 | 1.71 |  |
| 3 | Caecum | 23.75 | 26.13 | 2.38 | 5.21 |  |
| 4 | Colon | 23.23 | 26.05 | 2.82 | 7.07 | 4.63 |
| 5 | Colon | 24.11 | 26.09 | 1.98 | 3.95 |  |
| 6 | Colon | 25.12 | 26.64 | 1.52 | 2.87 |  |
| 7 | Duodenum | 25.13 | 26.63 | 1.50 | 2.84 | 2.16 |
| 8 | Duodenum | 25.67 | 26.56 | 0.89 | 1.86 |  |
| 9 | Duodenum | 25.37 | 26.21 | 0.84 | 1.79 |  |
| 10 | Ileum | 22.82 | 24.63 | 1.81 | 3.50 | 4.26 |
| 11 | Ileum | 25.11 | 26.92 | 1.81 | 3.51 |  |
| 12 | Ileum | 23.03 | 25.56 | 2.53 | 5.77 |  |
| 13 | Jejunum | 24.05 | 27.11 | 3.06 | 8.36 | 4.27 |
| 14 | Jejunum | 23.69 | 24.45 | 0.76 | 1.69 |  |
| 15 | Jejunum | 24.36 | 25.83 | 1.47 | 2.78 |  |

**Figure 2:**
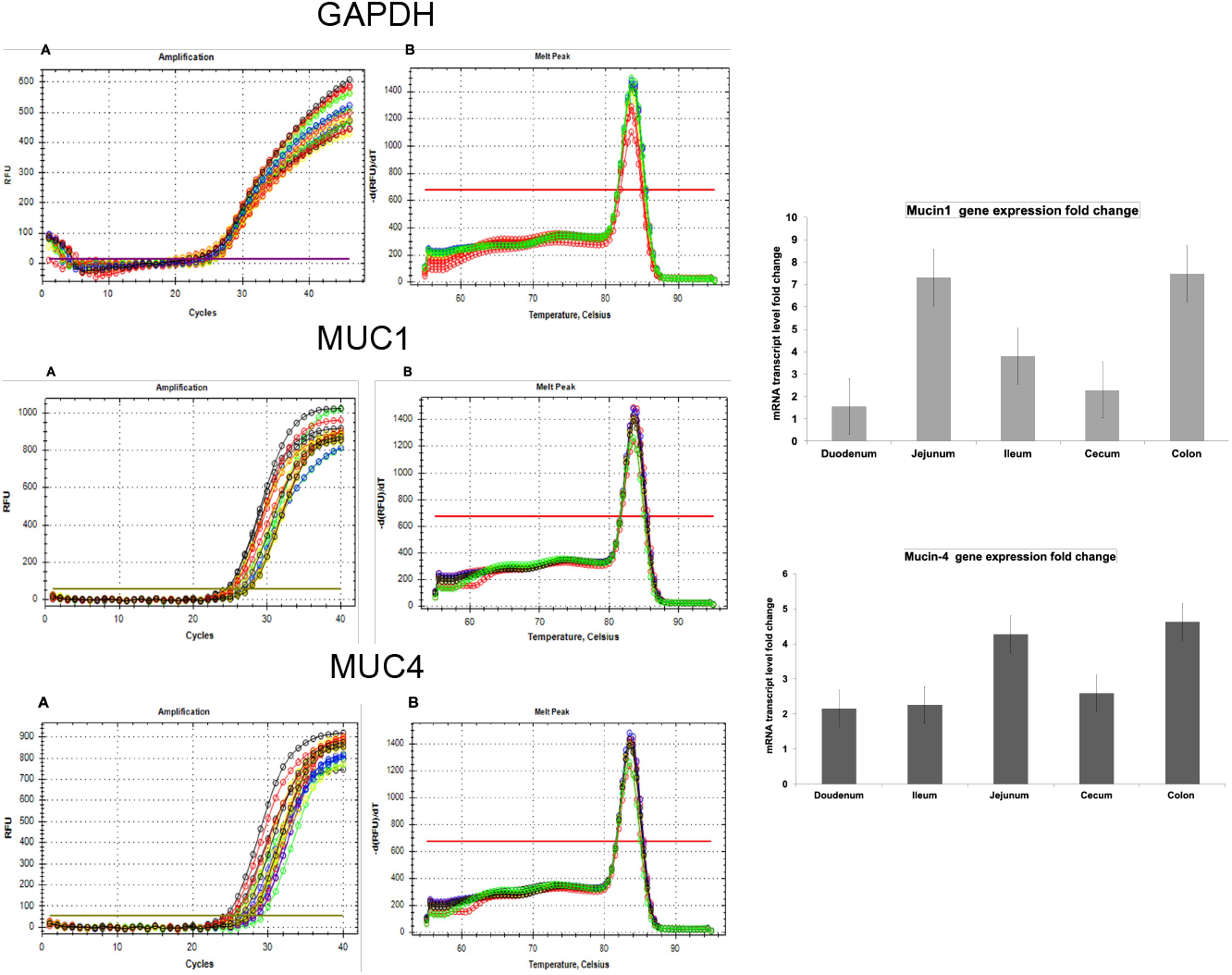
The RT-qPCR amplification plots of gene(A) and PCR melting peak (B) of the GAPDH MUC1 and MUC4 gene of the different regions of the small and large intestine of quail. The observable threshold cycles (Ct) numbers of expression were specified and different between regions of intestine are shown. The bar graph illustrates the level mRNA expressions of the MUC1 and MUC4 in the duodenum, jejunum, ileum, caecum and colon.

### Network analysis of quail mucins

20 quail specific mucins were identified in the Uniprot database, which are summarised in table 3. The major forms of mucins identified in quails were, MUC24, MUC15, MUC16, MUC6, MUC5, MUC4, MUC2, MUC3 and MUC1. Of these only five mucin types (MUC3, 5, 5BL, 6 and 15) showed network interactions in the STITCH database, which are summarised in figure 3. The major networks of these were predominantly mucin-mucin interactions in humans as well as quail (figure 3, table 4 and 5), suggesting preferentially homologous associations. Hence to assess which mucin combinations will have greater biochemical interactions, a detailed evaluation of intermolecular hydrogen bond formation between each mucin combinations was performed using the Chimera software. Of the 20 quail specific mucins only 15 mucin showed unique sequences for which either an AlphaFold (AF) or homology modelled (HM) 3D structure was available or could be generated. These 15 mucins were assessed for intermolecular hydrogen bond formation between them. The highest interactions were observed between the following combinations; MUC5A-MUC6, MUC5A-MUC5B, MUC5B-MUC6, MUC4-MUC6, MUC5A-MUC4 and MUC5B-MUC4. While MUC 15, 16 and 24 were observed to be least interacting with other mucin types (Figure 3). Our mucin-mucin network analysis indicated existence of two major categories of mucins i.e., 1) mucins (MUC 2, 4, 5 and 6) which form heterogenous networks and hence may be involved in blanketing the epithelium to provide protection and 2) mucins (MUC 1, 15, 16 and 24) which show poor affinity for interaction with other mucins and hence may be involved in forming only homogenous networks or functioning as solo in facilitating movement and transport of particles, bodies defence against pathogens, and various cellular processes and interactions including cell adhesion, differentiation, and inflammation.

**Table 3:** Various forms of mucins expressed in different breeds of quails.

| Uniprot ID | Gene | Mucin type | Species |
| --- | --- | --- | --- |
| A0A7K9Z0E4 | MUC15 | MUC15 | Odontophorus gujanensis (OG) |
| A0A8C2TUH6 | Mucin 15 | MUC15 | Coturnix japonica (CJ) |
| A0A7K9YXX0 | MUC16 | MUC16 | OG |
| A0A7K9YTP5 | Muc2_0 | MUC2 | OG |
| A0A8C2U6P4 | MUC2 | MUC2 | CJ |
| A0A226MFN5 | ASZ78_003200 | MUC2 | Callipepla squamata (CS) |
| A0A7K9YY45 | CD164 | MUC24 | OG |
| A0A7K9YS25 | Muc2l | MUC2l | OG |
| A0A8C2TP72 | MUC4 | MUC4 | CJ |
| A0A7K9ZAF7 | MUC4 | MUC4 | OG |
| A0A8C2TS61 | MUC4 | MUC4 | CJ |
| A0A7K9YT94 | Muc5ac_1 | MUC5A | OG |
| A0A7K9YR83 | MUC5B | MUC5B | OG |
| A0A8C2U825 | LOC107314549 | MUC5B | CJ |
| A0A226MF19 | ASZ78_003201 | MUC5B | CS |
| A0A8C2U8G1 | MUC6 | MUC6 | CJ |
| A0A7K9YTQ5 | Muc6 | MUC6 | OG |
| A0A226MUE7 | ASZ78_012728 | MUCL1 | CS |
| A0A8C2SVR3 | LOC107307211 | MUCL3B | CJ |
| A0A8C2U4E5 | LOC107314529 | MUCL5B | CJ |

**Table 4:**
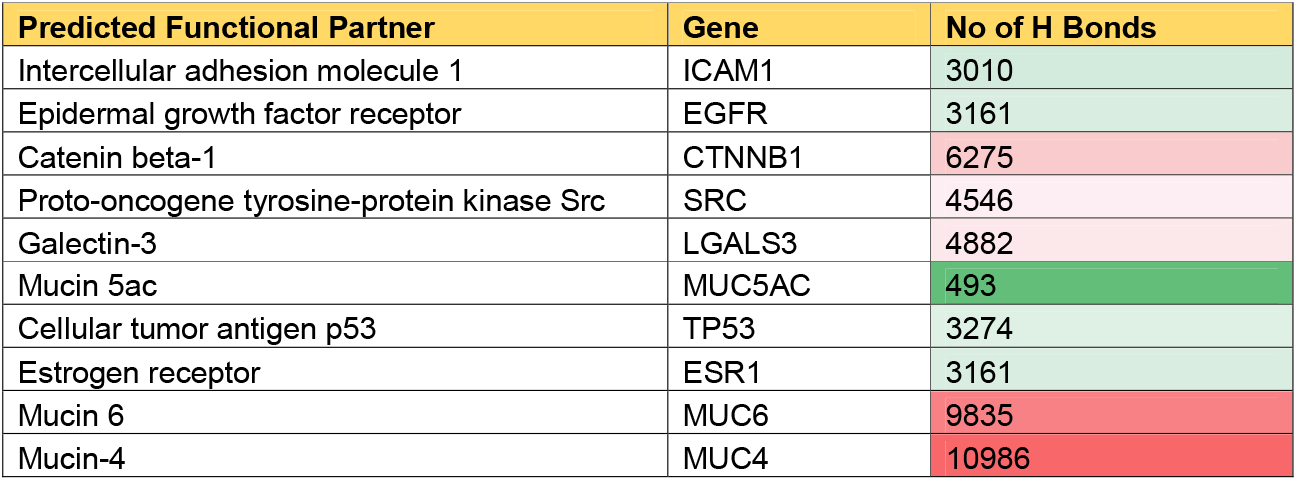
Network analysis of human MUC1.

**Table 5:** Network analysis of quail MUC4.

| Predicted Functional Partner | Gene | No of H Bonds |
| --- | --- | --- |
| Receptor tyrosine-protein kinase erbB-2 | ERBB2 | 9406 |
| MUC16 | MUC16 | 1993 |
| Mucin 6 | MUC6 | 15332 |
| MUC1 | MUC1 | 7874 |
| Mucin 2 | MUC20 | 6264 |
| MUC13 | MUC13 | 5239 |
| Mucin-7 | MUC7 | 2409 |
| Receptor tyrosine-protein kinase erbB-3 | ERBB3 | 9672 |
| MUC15 | MUC15 | 2755 |
| Mucin-21 | MUC21 | 10455 |

**Figure 3:**
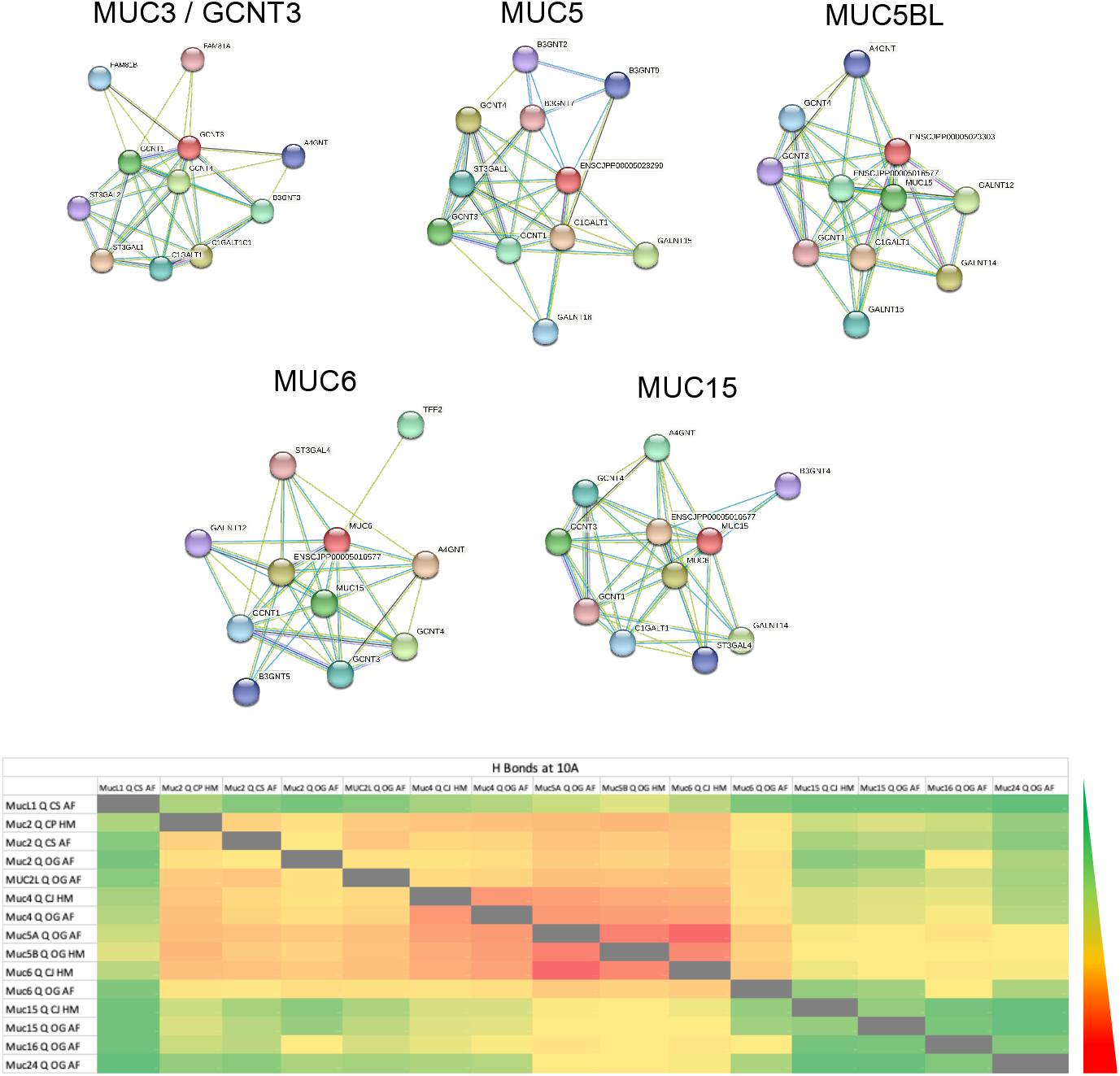
Network protein interaction of various quail mucin types in the STRING database. The heatmap represents the number of hydrogen bonds formed between each pair of quail mucins at 10 Armstrong evaluated in the Chimera software. 3D structure of 15 different quail specific mucins were used in this analysis. Abbreviations used: Odontophorus gujanensis (OG), Coturnix japonica (CJ), Callipepla squamata (CS), AlphaFold (AF), Homology Model (HM). Scale: Red to green represents high to low number of hydrogen bonds ranging from 25000 to 224 hydrogen bonds.

### Binding site analysis of quail mucins

Mucins play a critical role in protecting the epithelial lining of the gastrointestinal tract and perhaps various cell signalling process. Alterations in mucin production or the composition of the mucus layer is reported to occur in conditions like inflammatory bowel disease (IBD), Crohn’s disease, ulcerative colitis and colorectal cancers. Although currently drugs that directly target mucins are not available, considering the advancement in understanding of the biological role of mucins in various gastrointestinal pathophysiological process, it may merit development of drugs that selectively targets specific mucins. Hence in this study we analysed the binding sites of all the 15 quail specific mucins for which a 3D structure was available or could be generated (figure 4 and table 6). MUC5B and MUC6 had the most number of binding sites with high probability (> 0.8) scores. MUC 2, 4 and 5A although had more number of binding sites (>10 binding sites), their probability scores were lower (<0.6) suggesting their poor targetability or weaker interactions. MUC24 didn’t have any binding sites. All the other mucins (MUC 1, 15, and 16) which showed poor affinity interactions in our network analysis also had very few binding sites (<3 binding sites) with poor probability scores (<0.1). The sequence of the high affinity binding sites of MUC6 (A_1099 A_1102 A_1103 A_1106 A_1109 A_1110 A_1113 A_1121 A_1122 A_1127 A_1128 A_1129 A_1131 A_1156 A_1157 A_1158 A_1159 A_1160 A_1162 A_1172 A_1173 A_1174 A_1175 A_959 A_960 A_961 A_962 A_963 A_965 A_979 A_981 A_985 A_987), MUC5B (A_1123 A_1124 A_1127 A_1128 A_1136 A_1143 A_1146 A_1148 A_1150 A_1151 A_1156 A_1158 A_1161 A_1166 A_1167 A_1168 A_1177 A_1179 A_1182 A_1197 A_22 A_24 A_25 A_957 A_958 A_959 A_960 A_961 A_966 A_976 A_977 A_978) and MUC2 (A_1088 A_1089 A_1093 A_1101 A_1108 A_1110 A_1112 A_1115 A_1125 A_1126 A_1131 A_1132 A_1133 A_1142 A_1144 A_1147 A_921 A_922 A_923 A_924 A_925 A_941) identified here can be helpful in development of mucin specific selective small molecules or antibodies.

**Table 6:** Summary of the binding pockets in various quail mucins.

| **Mucin Type** | **Uniport ID** | **Number of Binding pockets (probability range)** |
| --- | --- | --- |
| MUC2_Q_CP_HM * | A0A8C2U6P4 | 30 (0.001 – 0.636) |
| MUC2_Q_CS_AF | A0A226MFN5 | 7 (0.004 – 0.082) |
| MUC2_Q_OG_AF | A0A7K9YTP5 | 5 (0.001 – 0.127) |
| MUC2L_Q_OG_AF | A0A7K9YS25 | 7 (0.003 – 0.208) |
| MUC4_Q_CJ_HM * | A0A8C2TS61 | 15 (0.002 – 0.319) |
| MUC4_Q_OG_AF | A0A7K9ZAF7 | 7 (0.021 – 0.44) |
| MUC5A_Q_OG_AF * | A0A7K9YT94 | 19 (0.001 – 0.295) |
| MUC5B_Q_OG_HM * | A0A7K9YR83 | 51 (0.005 – 0.855) |
| MUC6_Q_CJ_HM * | A0A8C2U8G1 | 34 (0.015 – 0.924) |
| MUC6_Q_OG_AF | A0A7K9YTQ5 | 4 (0.018 – 0.525) |
| MUC15_Q_CJ_HM | A0A8C2TUH6 | 2 (0.005 – 0.052) |
| MUC16_Q_OG_AF | A0A7K9YXX0 | 3 (0.017 – 0.098) |
| MUCL1_Q_CS_AF | A0A226MUE7 | 1 (0.028) |
| MUC24_Q_OG_AF | A0A7K9YY45 | None |
| MUC15_Q_OG_AF | A0A7K9Z0E4 | None |

Abbreviations used: *Odontophorus gujanensis* (OG), *Coturnix japonica* (CJ), *Callipepla squamata* (CS), AlphaFold (AF), Homology Model (HM).

**Figure 4:**
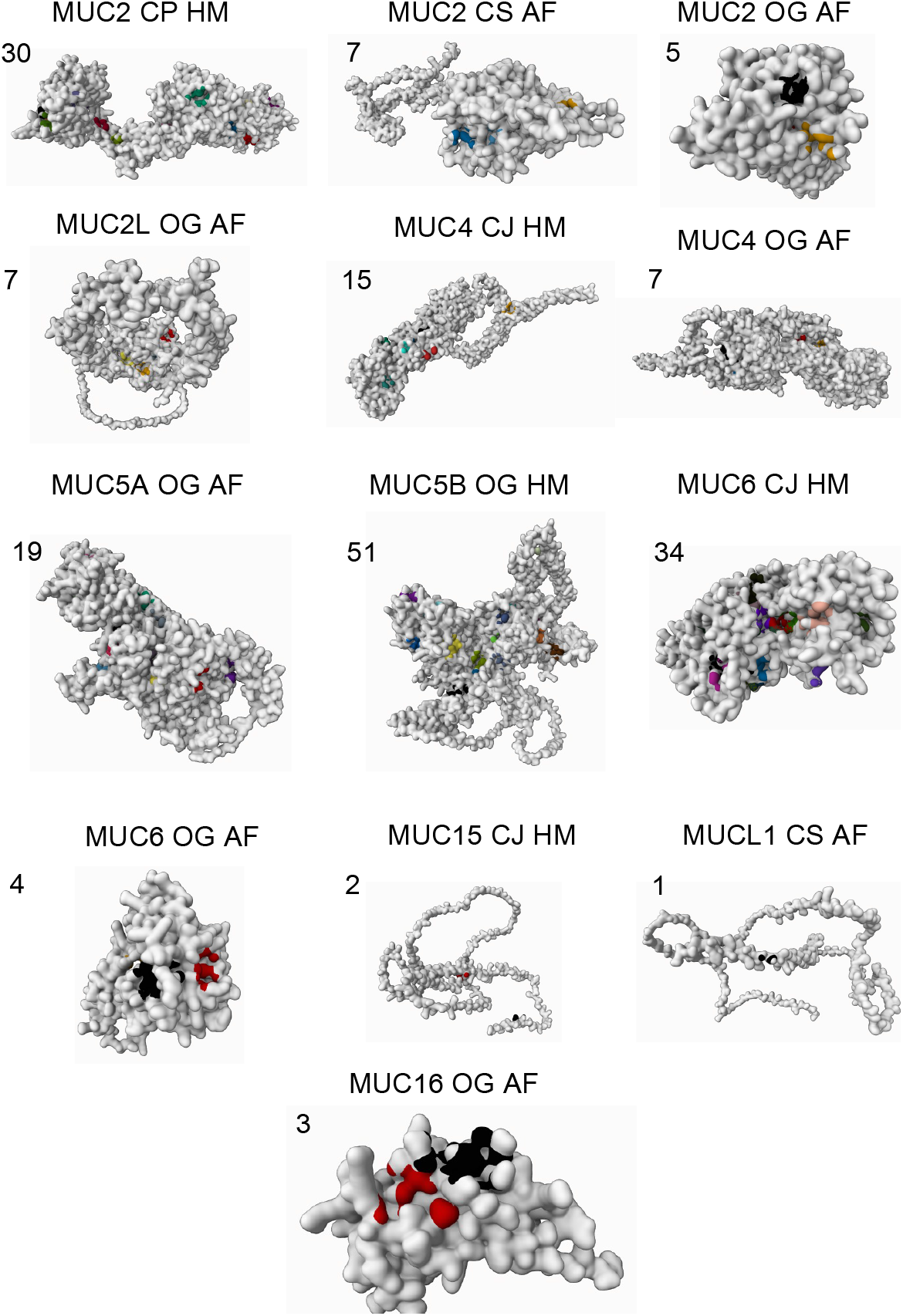
Binding site analysis of various quail mucin types. The values indicate the number of binding sites identified (represented in various colors). Abbreviations used: Odontophorus gujanensis (OG), Coturnix japonica (CJ), Callipepla squamata (CS), AlphaFold (AF), Homology Model (HM).

## Discussion

The findings of this study provide detailed insights into the histological features of intestines, mucin expression, mucin network interactions, and mucin binding site analysis of quails. Mucus play a vital role in maintaining the right environment of microflora of intestine, regulate nutrient transport of foods, and immune response in addition to preventing pathogen invasion[32, 33]. As mucins are the main components of the mucus[34, 35], in this study we investigated the various types of mucins reported in different breeds of quails and specifically evaluated the transcripts of two types of mucins (MUC1, and MUC4) in the various regions of quail intestines. Our study specifically identified two major categories of mucins based on their network interactions i.e., 1) high affinity heterogenous network mucins and 2) low affinity homogenous solo mucins, which in our view is vital to unravelling their role in various physiological processes and potential implications for gastrointestinal diseases.

Histologically, the quail intestines exhibited the typical layered structure consisting of mucosa, submucosa, muscularis, and serosa layers, which aligns with the existing knowledge from avian literature[36, 37]. The presence of these layers confirms the structural integrity and organization of the quail intestinal tract. Within the mucosa layer, three distinct layers of mucosa were observed consistently across all regions: a simple columnar epithelium with goblet cells on the basement membrane, an extended lamina propria with intestinal glands, and aggregations of lymphocytes surrounded by loose connective tissue[38, 39]. This histological assessment provides a comprehensive overview of the quail intestinal architecture and confirms its similarity to other avian species[40-42]. The staining analysis using alcian blue and PAS stains enabled the differentiation between acidic and neutral mucins. The predominance of blue-stained mucins with sporadic PAS staining indicates that the majority of mucins in the quail intestines are acidic mucins, while neutral mucins and glycogen content were minimally observed. The major expression of acidic mucin types in quail intestine sis consistent with a previous report[43]. Interestingly, there were regional variations in mucin expression, with higher expression of acidic mucins in the duodenum, ileum, caecum, and colon, and relatively higher expression of neutral mucins in the jejunum. This staining pattern suggests that acidic mucins play a prominent role in the protective functions of the quail intestinal mucosa, while neutral mucins may have specific functions in the jejunum. Also the relatively higher expression of acidic mucins in caecum, and colon justify the lubrication, protection and moisture retention role of this category of mucins which is facilitated by the high content of negatively charged sialic acid residues, higher glycosylation and large number of sulphate and carboxyl groups. The relatively higher expression of neutral mucins in the jejunum is a surprising observation in this study and the functional significance of this remains to be examined. Alterations in the balance between acidic and neutral mucins have been observed in various diseases[44, 45]. For example, an increase in acidic mucins and a decrease in neutral mucins have been reported in inflammatory conditions such as inflammatory bowel disease (IBD)[46-48]. In this context it remains to be seen if ratio of acidic to neural mucin (ANM ratio) can be developed and validated as a biomarker of IBD.

We identified 20 different types of mucin reported in various breeds of quails but focused our gene expression analysis using RT-qPCR on MUC1 and MUC4 transcripts in different regions of the quail intestine as these two are the major transmembrane mucins types which constitute intestinal mucosal layer[49, 50]. The detection of MUC1 and MUC4 mRNA in both the small and large intestine indicates their presence and potential involvement in various physiological processes. The jejunum and colon exhibited higher expression levels of both MUC1 and MUC4 mRNA compared to other regions. This suggests that these mucins may have specific roles in the jejunum and colon, potentially related to their functions in cellular processes, signalling, and protection. However, future studies will be helpful to look at the gene expression analysis of other types of mucins reported in this study, especially focusing on the results of our network analysis in which MUC 5 and 6 together with MUC 2 seem to have a major influence in mucin physiology in quails. The network analysis of quail mucins shed light on the interactions between mucins, providing insights into potential functional relationships and interplay. The identified mucin-mucin interactions, particularly among MUC2, MUC4, MUC5 and MUC6, suggest the existence of heterogeneous networks involving these mucins. These networks by forming a protective barrier blanket the epithelium, and provide defence against pathogens. On the other hand, mucins such as MUC1, MUC15, MUC16, and MUC24 showed poor affinity for interactions, potentially indicating their involvement in individual functions or cellular processes. Also it remains to be established if biomarkers based on the variations in the ratio of heterogeneous to homogenous mucins could be biologically relevant.

The heterogeneous mucin networks are characterized by interactions between different mucin types and other components of the mucus layer[51, 52]. The diversity of mucin species within the network allows for the formation of a complex matrix that helps blanket the epithelium and provides a physical barrier against pathogens, toxins, and mechanical stress[53]. The interplay between different mucins within these networks perhaps contributes to mucus viscosity, lubrication, and the ability to trap and remove foreign particles[54]. Heterogeneous mucin networks are hence crucial for mucosal defence and play a significant role in preventing infections and maintaining tissue homeostasis[55]. In contrast the homogenous mucins exhibit poor affinity for interactions with other mucins and are more limited in their ability to form complex matrices. Instead, they may function as individual entities or interact primarily with specific molecules or cell surface receptors. Homogeneous mucin networks may be involved in facilitating movement and transport of particles, as well as participating in various cellular processes and interactions, including cell adhesion, differentiation, and inflammation. Their relatively lower affinity for interactions might allow them to act more independently, flexibly or in a targeted manner, potentially influencing specific cell signalling pathways or functions. Further understanding of the heterogenous versus homogenous mucins will require their selective targeting and to the best of our knowledge such selective targeting tools are currently lacking. Hence as a first step towards this selective targeting process, we have identified the specific binding sites of the 15 quail specific mucins for which the 3D structures were generated in this study.

The binding site analysis of quail mucins aimed to identify potential targets for selective drugs or therapeutic interventions. Among the quail-specific mucins, MUC5B and MUC6 exhibited the highest number of binding sites with high probability scores, indicating their potential as targets for drug development. MUC2, MUC4, and MUC5A showed a larger number of binding sites, albeit with lower probability scores, suggesting that their targetability might be more challenging. MUC24 displayed no binding sites, while other mucins (MUC1, MUC15, and MUC16) exhibited few binding sites with poor probability scores. We report in this study a high affinity binding site sequence of MUC2, MUC5B and MUC6, which will be valuable in developing selective small molecules or antibodies against the binding site. MUC1 and MUC 4 are reported to be involved in several important functions in the digestive system, including lubrication of the digestive tract, protection of the epithelial surface from mechanical and chemical damage, and regulation of the immune response[56-58]. However, in contrast to these reports, the network analysis from this study suggests a major role for MUC5B MUC6 and MUC2 in quails, the pathophysiological significance of which remains to be examined in future studies. While the role of MUC4 in quail intestines cannot be excluded considering its higher number of low affinity binding sites, the role for MUC1 in quails seems to be less relevant considering identification of only one very low affinity binding site.

In conclusion, this study provides valuable insights into the composition, expression, network interactions, and potential targetability of mucins in the quail intestine. Understanding the role of mucins in gastrointestinal physiology and their implications for gastrointestinal diseases is crucial for advancing our knowledge of mucosal protection, cellular processes, and therapeutic strategies. The findings presented here lay the foundation for further research and potentially guide the development of targeted interventions that modulate mucin functions in avian species and potentially in humans as well.

